# Growth potential of *Listeria monocytogenes* in twelve different types of RTE salads: impact of food matrix, storage temperature and shelf life

**DOI:** 10.1101/261552

**Authors:** Matthias Ziegler, David Kent, Roger Stephan, Claudia Guldimann

## Abstract

Listeriosis is a food borne disease associated with high hospitalization and fatality rates; in 2014, EU member states reported 2194 cases with 98.9% hospitalization rates and 210 fatalities. Proper risk analysis and the development of effective food safety strategies critically depend on the knowledge of the growth characteristics of *L. monocytogenes* on the product in question. Ready-to-eat (RTE) salads present a challenge in this context due to the absence of a heat treatment step before consumption and the interaction of pathogens with the plant microbial microbiota. This study provides challenge-test based data of the growth characteristics of *L. monocytogenes* on twelve RTE salads. The food matrix, storage time and storage temperature were factors with a significant impact on the growth of *L. monocytogenes*. While most tested salads permitted a significant increase of *L. monocytogenes* in at least one of the tested conditions, no growth was observed on celeriac, carrot and corn salad products. There was a considerable increase in growth at 8 °C compared to 5 °C. Our data indicate that the reduction of the storage temperature at retail level to 5 °C and product shelf life could help mitigate the risk of *L. monocytogenes* in RTE salads.

## 1. Introduction

Listeriosis remains one of the most severe food borne diseases. The relatively low incidence (0.46 cases / 100’000 in the EU in 2015 with an ongoing, significant increase since 2008) (European Food Safety Authority European Centre for Disease Prevention and Control, 2016) is offset by the a high mortality rate (15-30 deaths / 100 cases) (Barton Behravesh et al., 2011; de Valk et al., 2005; Popovic, Heron, & Covacin, 2014; WERBER et al., 2012), mostly due to severe forms of Listeriosis like central nervous system infections, septicemia and abortions/neonatal infections (Allerberger & Wagner, 2010). Food borne outbreaks of Listeriosis have been associated in the past with dairy products, fish and seafood, meat products, fresh fruit and vegetables, and ready-to-eat (RTE) products (Datta, Laksanalamai, & Solomotis, 2013; European Food Safety Authority European Centre for Disease Prevention and Control, 2016). Within the RTE food category, raw products without a heat-treatment step during the production process (e.g. RTE-salads, fruit, vegetable or dairy products) have an inherently increased risk for contamination with pathogens. *Listeria monocytogenes* is a challenging problem in this context and a priority for food producers due to its ubiquitous presence in the environment and the ability to grow at refrigeration temperatures. A current literature review of studies on the contamination levels of RTE salads with mixed ingredients (defined as raw salads combined with processed foods such as ham, chicken, salmon or pasta) concludes that within the EU, about 2-10% of products were contaminated with *L. monocytogenes* (Söderqvist, 2017). In 2015 the EU member states reported 0.04% of tested RTE salad products to exceed the legal limit of *L. monocytogenes* (European Food Safety Authority European Centre for Disease Prevention and Control, 2016). In Switzerland, leafy green RTE salads caused an outbreak in 2013-2014 (Stephan et al., 2015). Producers of RTE salads are faced with two fundamental and contradictory requirements: (i) to provide food at the highest possible safety standards while (ii) meeting the demand from retailers and consumers for food with an increasingly long shelf life. The EU food safety regulations limit *L. monocytogenes* to < 100 colony forming units (CFU)/g at the end of the shelf life of a food product. Accordingly, different food safety criteria are applicable based on whether a food supports the growth of *L. monocytogenes*. For food that permits growth of *L. monocytogenes*, the producer must test for absence in 25 g at the time the food leaves the immediate control of the producer, while for food that does not permit growth of *L. monocytogenes* the food safety criterion is < 100 CFU/g at the end of the shelf life (EC regulation No 2073/2005). To assess the risk associated with extending the shelf life, it is therefore crucial to determine the growth potential of *L. monocytogenes* in the respective food product.

The aim of this study was to determine the growth potential of *L. monocytogenes* on twelve RTE salad products, under packaging and storage conditions that mirror retail and home conditions.

## 2. Materials and methods

All experiments were carried out in three independent replicates.

### 2.1. Bacterial strains, growth conditions and subtyping

The three strains of *L. monocytogenes* used in this study were all isolated from a RTE salad production facility (table 1). Stock cultures of *L. monocytogenes* were maintained at −80 °C in brain heart infusion (BHI; Oxoid, Basel, Switzerland) broth with 15% glycerol. To prepare the inocula, stock cultures were streaked on BHI agar plates and incubated overnight. A single colony was inoculated into 5 ml BHI broth and incubated overnight (37 °C, 200 rpm), subcultured in the morning 1:100 into 5 ml fresh BHI broth, and incubated for 6 h (37 °C, 200 rpm) to obtain an early-stationary-phase culture (9.6 ± 0.1 log CFU/ml). This culture was then incubated at 5 °C for 20 h for cold adaptation. Strain pools were obtained by combining equal quantities of the cold adapted stationary phase cultures.

**Table 1.**

| Species | Strain designation | Serotype | ST | CC | Lineage |
| --- | --- | --- | --- | --- | --- |
| *Listeria monocytogenes* | N16-1125 | 1/2a | ST91 | CC14 | II |
| *Listeria monocytogenes* | N16-0716 | 1/2b | ST517 | CC517 | I |
| *Listeria monocytogenes* | N16-0855 | 4b | ST6 | CC6 | I |

ST: sequence type. CC: clonal complex

The three strains used in this study were serotyped using listeria antisera from Denka Seiken (Pharma Consulting, Burgdorf, Switzerland) according to the manufacturer’s protocol. They were then further characterized using multi locus sequence typing (MLST) based on seven housekeeping genes as described by Ragon et al. (Ragon et al., 2008) with minor modifications. DNA was extracted using the DNeasy Blood&Tissue Kit (Qiagen, Hilden, Germany) according to manufacturer’s protocol. The PCRs were performed using a HotStarTaq master mix (Qiagen, Hilden, Germany) and the following conditions: for *bglA, cat* and *ldh*: 95 °C for 15 min, followed by 41 cycles of 95 °C for 30 s, 45 °C for 30 s and 72 °C for 30 s; for *abcZ, dapE, dat*, and *lhkA*: 95 °C for 15 min followed by 35 cycles of 95 °C for 30 s, 52 °C for 30 s, and 72 °C for 30 s. The final extension was at 72 °C for 10 min for all amplifications.

PCR products were purified using the GenElute PCR Clean-Up Kit (Sigma, Steinheim Germany) according to manufacturer’s protocol and were sequenced using primers oR and oF (Ragon et al., 2008). The clonal complex (CC) and sequence type (ST) were determined using the *Listeria* MLST database hosted at the Institute Pasteur (http://bigsdb.pasteur.fr).

### 2.2. Cold growth in rich medium

Growth at cold storage temperatures was determined for the three *L. monocytogenes* strains used in this study individually in a control experiment. Early stationary phase cultures were obtained and cold adapted as described above. This culture was then diluted 1:1000 into 10 ml fresh BHI broth and incubated at 5 °C and 8 °C for 11 days. CFU/ml were determined by plate counting 10 µl aliquots from each culture at t=0 and after 1, 4, 5, 6, 7, 8, and 11 days. Aliquots were serially diluted in Maximum Recovery Diluent (MRD; Oxoid, Basel, Switzerland) and 10 µl of the serial dilutions were spotted on BHI agar plates and run over by tilting as described by Kuehbachler et al. (Kühbacher, Cossart, & Pizarro-Cerdá, 2014). Plates were incubated at 37 °C for 24 h. Final results were expressed as log CFU/ml.

### 2.3. Salad products

The range of salads was limited to plant based products without added ingredients of animal origin, and were chosen to represent commonly sold RTE salad products. A total of 12 different RTE salad products from a producer in Switzerland were used for the challenge tests (table 2). Three RTE salad products were packaged under modified atmosphere. To avoid sampling effects due to uneven distribution of *L. monocytogenes* within a package, the whole content of a bag was used for analysis. For this purpose, 30 g salad portions (parsley: 20 g portions) were produced specifically for this study, shipped to our facility under preservation of the cold chain at 5 °C and inoculated 12-24 h after production. The packaging foil and, where appropriate, the modified atmosphere was identical to the larger packages that were produced for retail.

**Table 2:**
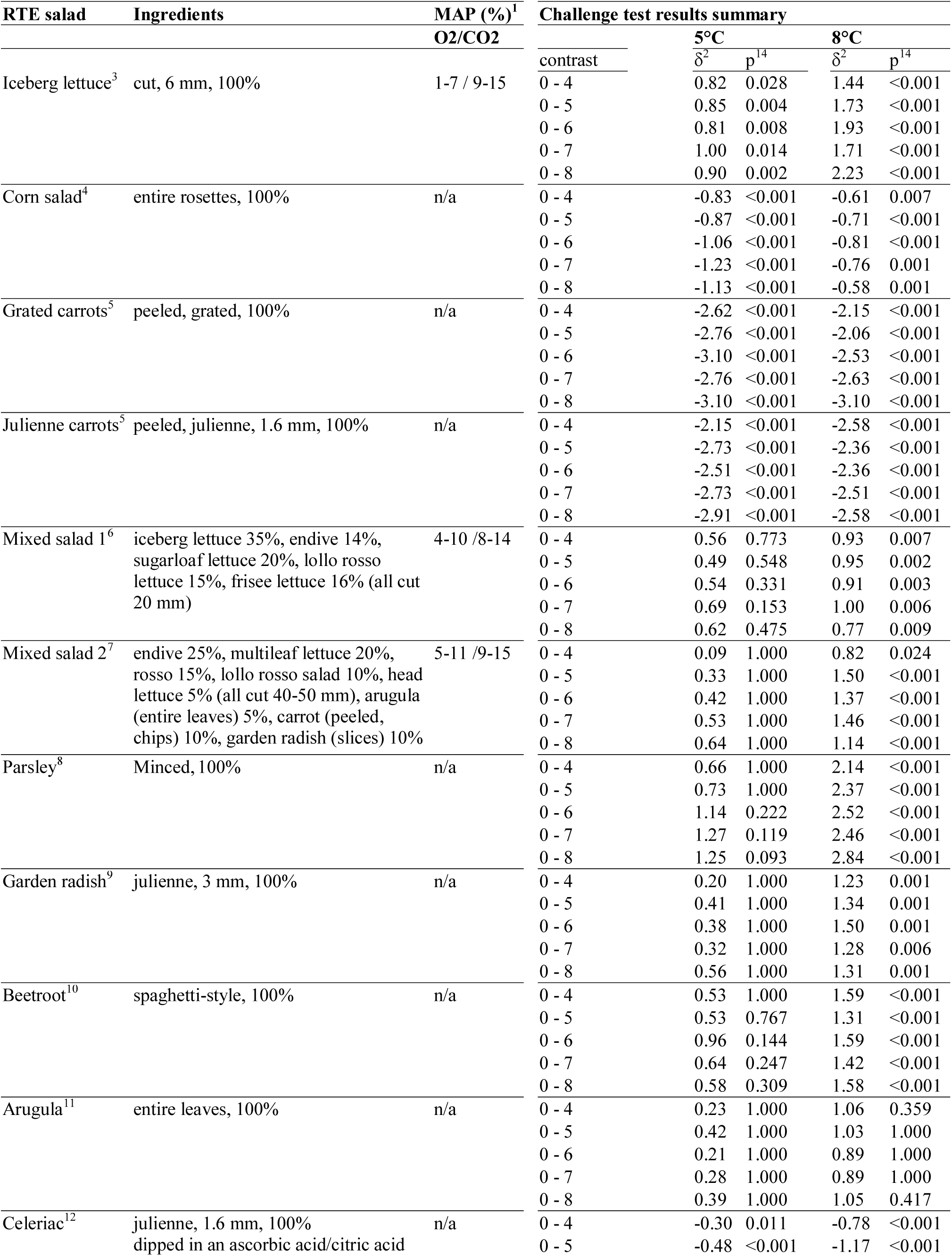
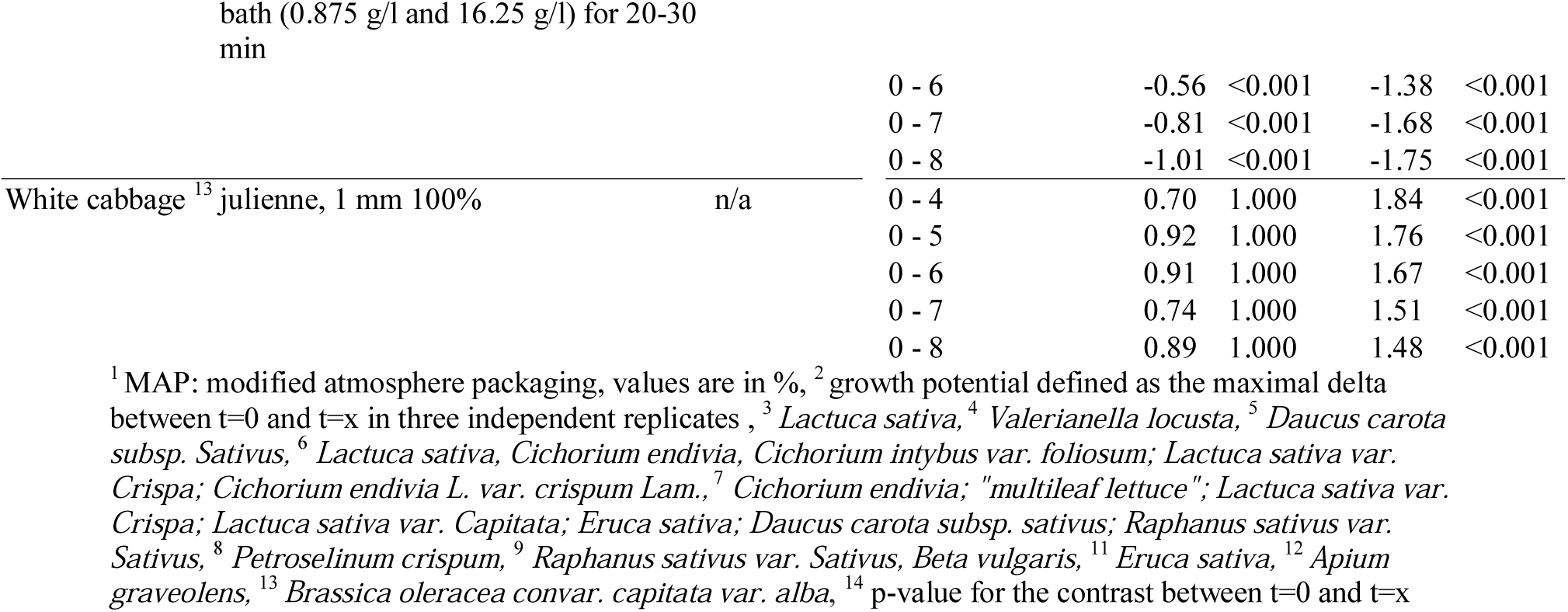

### 2.4. Inoculation of the food matrix

To ensure that all three strains used in this study had a similar growth phenotype on salad products, a first experiment was performed with the three *L. monocytogenes* strains inoculated individually on two RTE salad products (iceberg lettuce and arugula). Based on the results of these preliminary experiments, the other ten RTE salad products were inoculated with an equalized pool of the three strains.

To achieve a final bacterial load of 5 log CFU/g in the products, the cold adapted stationary phase culture was serially diluted in MRD and 1 ml of the appropriate dilution was homogeneously distributed over the product. This relatively high concentration was chosen to be able to accurately quantify not only an increase, but also a decrease in CFU/g. Unlike in other products, the normal microbiota in RTE salad products is in the range of 5-6 log CFU/g and the administered inoculum is therefore only a fraction of the total bacteria present in the samples. The inoculum was administered through a septum of scotch tape using a syringe and a gauge 22 needle. Immediately after inoculation, the syringe hole was sealed with a second scotch tape to maintain the modified atmosphere. Negative control samples were inoculated in the same way with 1 ml MRD, and for RTE salad products packaged under modified atmosphere, uninoculated bags of product served as controls for the modified atmosphere. After inoculation, all samples were shaken for 1 min in a standardized manner to optimize the distribution of the inocula. To preserve the cold chain, the bags of salad and the bacteria were kept on ice at all times during these procedures.

### 2.5. Storage conditions

The RTE salad products were stored at 5 °C and 8 °C, for 4, 5, 6, 7 and 8 days. Temperature in both cold rooms was continuously controlled and recorded with temperature loggers (EasyLog, Lascar Electronics, Pennsylvania, USA).

### 2.6. Microbiological analyses

Enumeration of *L. monocytogenes*, Enterobacteriaceae and the total viable count (TVC) were performed according to ISO 11290-2:1998, ISO 21528-2:2004 and ISO 4833-2:2013 respectively with minor modifications.

*L. monocytogenes* and TVC were determined immediately after inoculation (t=0) and 4, 5, 6, 7 and 8 days after inoculation. The counts of Enterobacteriaceae were determined at t=0 and 8 days after inoculation. At each time point, one inoculated sample and one negative (uninoculated) control sample per temperature were analyzed. The whole content of a unit was transferred into sterile stomacher bags, diluted 1:10 with MRD and homogenized for 30 s in a Stomacher® 400 Circulator (Seward, Worthing, United Kingdom). For products packaged under modified atmosphere, the gas composition in the bag was measured before opening and compared to gas composition in uninoculated packages using a gas detector (CheckPoint, Dansensor, Neuwied, Germany). Serial dilutions in MRD were prepared and 0.1 ml was spread-plated on the following agar plates in duplicate: PALCAM (Merck, Darmstadt, Germany) for the enumeration of *L. monocytogenes* (aerobic incubation for 24 h at 37 °C); plate count (PC; Oxoid, Basel, Switzerland) for TVC (aerobic incubation for 48 h at 37 °C); violet red bile glucose (VRBG; BD, Allschwil Switzerland and Merck, Darmstadt Germany) for Enterobacteriaceae (anaerobic incubation for 48 h at 37 °C). The average of the duplicate plates was calculated and expressed as log CFU/g. The limit of detection was 2 log CFU/g.

### 2.7. Calculation of the growth potential δ

For each time point at each temperature, the difference between the log CFU/g at the evaluation point and the log CFU/g at the beginning of the challenge test was calculated for each of the three independent replicates. The growth potential δ was defined as the highest value obtained among three replicates. When δ was higher than 0.5 log CFU/g the RTE salad product was classified as “able to support the growth of *L. monocytogenes*” at the corresponding temperature. If δ was ≤ 0.5 log CFU/g the RTE salad product was classified as “unable to support the growth of *L. monocytogenes*”.

### 2.8. Statistical analysis

Statistical analysis and graphics were performed in R (Version 3.4.0) (R Core Team, 2015) using R studio (Version Version 1.0.143) (RStudio Team, 2015). A linear mixed effects model was calculated using lmer in LmerTest (Kuznetsova, Bruun Brockhoff, & Bojesen Christensen, 2016) with up to three-way interactions between time, salad and temperature as random effects and replicate as fixed effect. Pairwise contrasts were calculated using lsmeans (Lenth, 2016). The ggplot2 package (Wickham, 2009) was used for visualization. Holm-Bonferroni adjusted p-values were calculated for multiple comparisons within one type of salad.

To determine if the three *L. monocytogenes* strains differed from each other in their growth on salad, linear mixed effects models were calculated; model selection was AIC based. An ANOVA was used to determine if the factor “strain” had an effect on the outcome “log CFU/ml”.

The R scripts can be downloaded from (insert link R_scripts_and_data).

## 3. Results and Discussion

### 3.1. Strain characterization and results from the proof of concept experiments

The three strains used in this study belonged to serotype 1/2a, 1/2b and 4b, CC 14, 517 and 6 and ST 91, 517 and 6 respectively (table 1).

A series of experiments was performed to ensure the soundness of the experimental setup. The cold-growth ability of the strains used in this study was confirmed in rich medium. All three *L. monocytogenes* strains were able to grow at the 5 °C and 8 °C in rich medium and reached early stationary phase after eleven days at 5 °C vs. after five days at 8 °C. There was no difference in cold growth between the three strains (supplementary figure 1).

To avoid bias caused by one of the strains used in the pool outgrowing the others, the first experiments (on iceberg lettuce and arugula samples) were performed with each strain individually. Since there was no significant effect of “strain” on “log CFU/ml” in the outcome (F=2.57, p=0.08), all further experiments were performed using a pool of the three strains. For the final analysis, the results from the individual strains on iceberg lettuce and arugula were combined to be comparable to the results from the experiments with the pooled strains.

The temperature was logged hourly for the 5 °C storage unit (mean 4.9 °C, SD 1.13, maximum 8 °C) and the 8 °C storage unit (mean 8.11 °C, SD 0.53, maximum 13 °C) over the complete duration of the experiments (152 days).

For those salads packaged under modified atmosphere, the gas composition was monitored. In mixed salad 1, none of the values were outside the range specified by the producer, while in the iceberg lettuce, some of the units stored at 8 °C were out of range. Many of the mixed salad 2 units were out of range, including some of the uninoculated controls (supplementary table 1). This may indicate a partial failure of this specific packaging/produce combination to reach a steady state between plant respiration and diffusion of gases through the packaging foil. The airtightness of the scotch tape used to seal the inoculation hole was tested by puncturing three bags and applying the seal before storing them for 72 h with 2.2 kg weight on top of them. None of the bags deflated (data not shown).

### 3.2. Growth of *L. monocytogenes* on RTE salad products

We found that storage temperature, storage duration and the food matrix with its associated microbiota had a significant impact on the growth potential of *L. monocytogenes* under the tested conditions.

#### Impact of temperature on the growth potential of L. monocytogenes

At 5 °C, the growth potential of *L. monocytogenes* exceeded 0.5 log CFU in both mixed salads, parsley, iceberg lettuce, garden radish, beetroot and white cabbage at least on one time point (figure 1, table 2). This was either at very late time points (e.g. at t=8 for garden radish) and/or not by a large margin, e.g. the growth potential was between 0.5-1 log CFU. Also, the increase in CFU/g between t=0 and the later time points was only statistically significant in iceberg lettuce (p < 0.05, table 2). Accordingly, at 5 °C the risk to exceed the food safety criterion of < 100 CFU/g for *L. monocytogenes* during shelf life could be contained for most salads. If the limit of detection was to be lowered to 10 CFU/g at the routine testing level, a growth potential of < 1 log CFU over the shelf live would still contain the bacterial load with *L. monocytogenes* to below 100 CFU/g and therefore be considered tolerable for foods not intended for infants or other vulnerable groups such as hospital patients.

**Figure 1:**
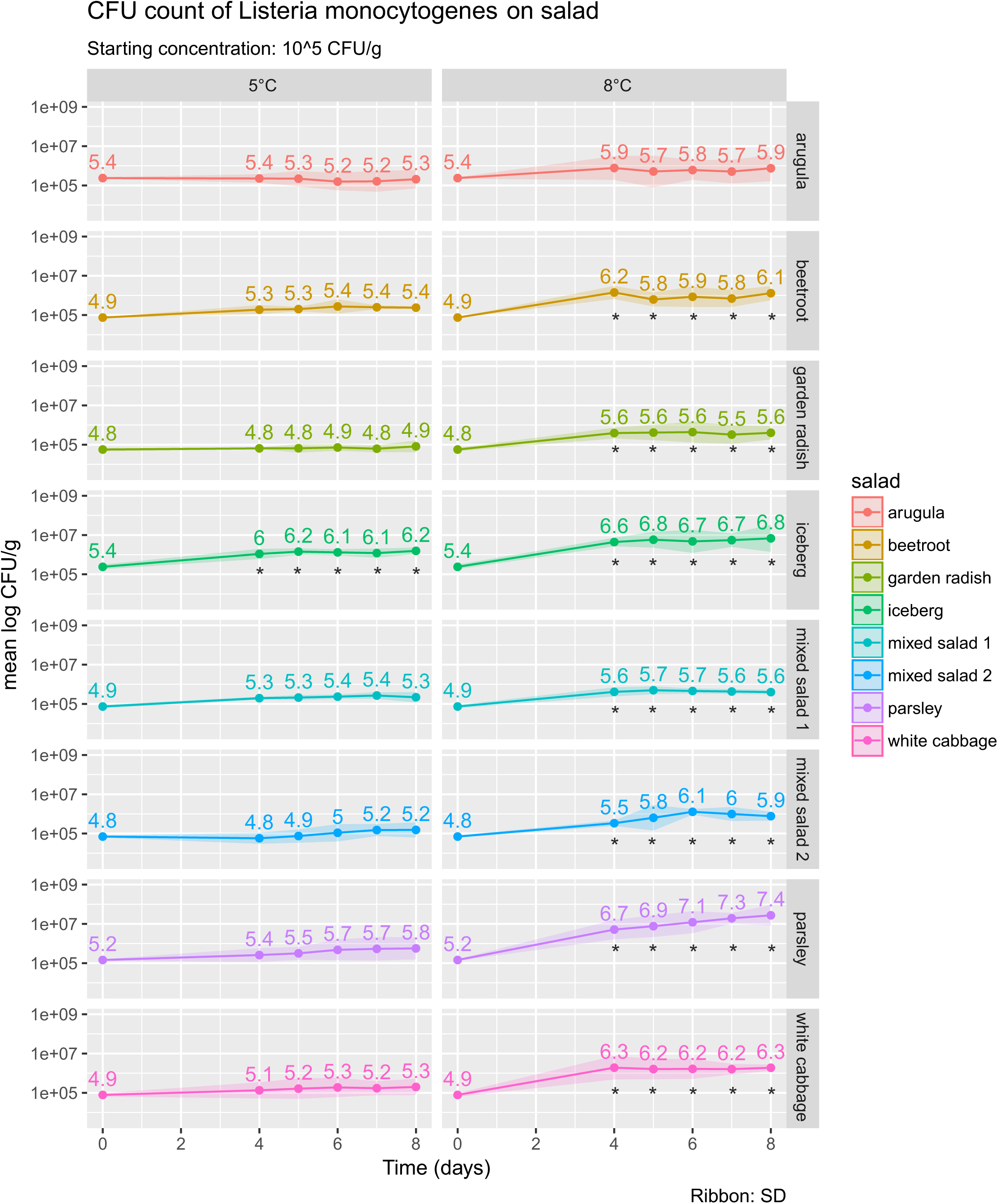
*L. monocytogenes* on RTE salads that supported growth. The numbers above time points reflect mean log CFU/g. Asterisks denote time points where the log CFU/g were significantly different from t=0 (p < 0.05). The ribbon around the line represents the standard deviation.

At 8 °C, the situation was drastically different. With the exception of corn salad, celeriac and carrots, *L. monocytogenes* exhibited a growth potential over or very close to 1.0 CFU. This increase in CFU/g between t=0 and the later time points was statistically significant in all salads except arugula (p < 0.05, table 2). In this situation, it is crucial to test at the stricter food safety criterion of “absent in 25 g” before the food leaves the immediate control of the producer.

The strikingly higher growth potential of *L. monocytogenes* at 8 °C compared to 5 °C may also challenge the common practice to store RTE products at 8 °C at retail level. Our data suggest that in some cases, growth of *L. monocytogenes* to > 100 CFU/g during shelf life could be mitigated by a stricter temperature management at retail level to limit storage temperature of RTE salad products to a maximum of 5 °C.

#### Impact of time on the growth potential of L. monocytogenes

As expected, time was a crucial factor in the outcome of total counts (CFU/g) of *L. monocytogenes*. In many of the salad products that supported the growth of *L. monocytogenes* (e.g. garden radish and white cabbage at 8 °C and iceberg lettuce at 5 °C and 8 °C), *L. monocytogenes* increased significantly between t=0 and t=4, after which a steady state was reached where no more significant increase occurred. A potential explanation for this plateau might be the interaction with the plant microbiota associated with individual plant species that limits further growth. In contrast, parsley at 8 °C permitted a significant increase in CFU/g also between t=4 and the later time points. This suggests that minced parsley is a high-risk product that supports substantial growth of *L. monocytogenes*.

#### Impact of the food matrix and its associated microbiota on the growth potential of L. monocytogenes

The food matrix had an impressive impact on the growth potential of *L. monocytogenes*. Four of the twelve RTE salad products tested in this study did not support the growth of *L. monocytogenes* in a significant way under the tested conditions, while the other eight products supported growth of *L. monocytogenes* at varying levels (table 2). Potential reasons for these vast differences may lie in the normal microbiota of the plant, the composition of the product, the level of processing of the plant matrix (e.g. availability of plant juice depending on the level of cutting vs. whole leaves), and in the natural defense mechanisms of the plants in question.

The product group that supported growth of *L. monocytogenes* comprised iceberg lettuce, parsley, arugula, beetroot, garden radish, white cabbage and the mixed salads. The maximal growth potential of 2.84 log CFU was observed in parsley (t=8, 8 °C) (figure 1).

*Listeria* has previously been shown to grow on iceberg lettuce (at 3, 5 and 10 °C) (Beuchat & Brackett, 1990b; Koseki & Isobe, 2005), arugula (6 days at 7 °C) (Sant’Ana, Barbosa, Destro, Landgraf, & Franco, 2012) and “green salad” (6 days at 7 °C) (Sant’Ana et al., 2012). Conflicting data exist on raw cabbage: one study on white cabbage found considerable growth (15 days at 4 °C, 7.5 days at 10 °C) (Wang et al., 2013), while another study on RTE cabbage found a reduction in *L. monocytogenes* numbers (6 days at 7 °C = −0.21 (SD 0.9) log CFU/g) (Sant’Ana et al., 2012). However, it is unclear if the cabbage in this latter study was processed or raw. Juice from fermented cabbage might have some antimicrobial properties against Gram negative organisms (Gogo, Shitandi, Lokuruka, & Sang, 2010).

Celeriac, carrots and corn salad did not support the growth of *L. monocytogenes*. At no point in time did the growth potential of *L. monocytogenes* exceed 0.5 log CFU in these products. In fact, the CFU/g *L. monocytogenes* dropped significantly between t=0 and the later time points at both 5 and 8 °C (figure 2). To our knowledge, this is the first report on the fate of *L. monocytogenes* on corn salad and celeriac. In contrast to the celeriac root, temperature-dependent growth of *L. monocytogenes* has been shown on celery stalks (decrease in CFU/g over 7 days at 4 °C (Vandamm, Li, Harris, Schaffner, & Danyluk, 2013) vs. growth or minimal increase at 10 °C over 7 days (Kaminski, Davidson, & Ryser, 2014; Vandamm et al., 2013). Also, celery stalks have been implied in an outbreak of listeriosis (Gaul et al., 2013). The relatively high content of nitrite in celery in combination with the fine cut of the celeriac julienne analyzed in the present study might provide a possible explanation for our observations; in fact celery powder is used as a natural source of nitrite in meat curing processes for its antimicrobial activity and coloring properties (Buchanan, Stahl, & Whiting, 1989; Junttila, Hirn, Hill, & Nurmi, 2016; McClure, Kelly, & Roberts, 1991). The anti-listerial effect of carrots has been attributed to phytoalexins produced by carrots in response to fungal infections and other types of stress (Abdul-Rauf, Beuchat, & Ammar, 1993; Babic, Nguyen-the, Amiot, & Aubert, 2008; Beuchat & Brackett, 1990a; Kurosaki & Nishi, 1983). It is conceivable that harvesting might result in increased concentrations of phytoalexins in carrots due to microlesions in the plants. The activity of phytoalexins is mainly directed against fungi and Gram+ organisms (Kurosaki & Nishi, 1983). This would spare the mostly Gram-natural microbiota of the plants, which is reflected in the fact that the total viable count increased over time in all salads except celeriac at 5 °C.

**Figure 2:**
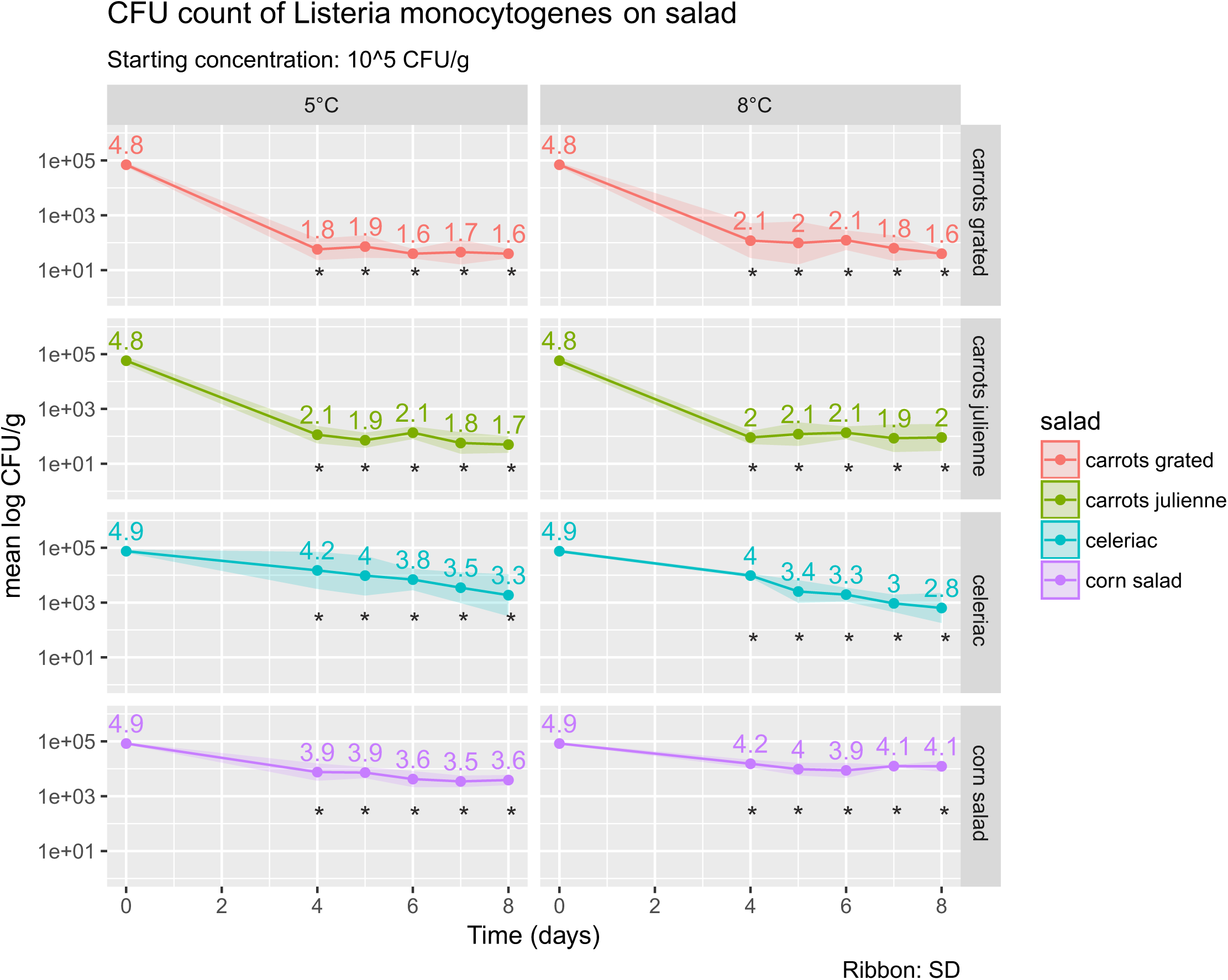
*L. monocytogenes* on RTE salads that did not support growth. The numbers above time points reflect mean log CFU/g. Asterisks denote time points where the log CFU/g were significantly different from t=0 (p < 0.05). The ribbon around the line represents the standard deviation.

### 3.3. Total viable count on RTE salad products

The initial total viable count ranged from 4.9 log CFU/g (celeriac (SD=0.09), white cabbage (SD=0.14)) to 6.7-6.8 log CFU/g (parsley (SD=0.44), arugula (SD=0.71), garden radish (SD=0.24)). At 8 °C, all products showed a significant increase in TVC at all time points, reaching as much as 9 log CFU/g (parsley). At 5 °C, celeriac (all time points) and arugula (all time points except at t=8) showed no significant increase in TVC. In all other products, there was a significant increase in TVC (figure 3). While other authors found similar counts for the TVC on iceberg lettuce (5.61 ± 0.41 log CFU/g) (Koseki & Isobe, 2005) and carrots (5.79 ± 0.04 log CFU/g) (Sant’Ana et al., 2012), studies found higher numbers for cabbage (7.67 ± 0.09 log CFU/g) (Sant’Ana et al., 2012), arugula (8.11 ± 0.16 log CFU/g) (Sant’Ana et al., 2012) and corn salad (6.63-6.85 log CFU/g) (Wei, Wolf, & Hammes, 2005). However, the validity of comparisons between studies is at least questionable, as many factors like plant variety and maturity, the processing conditions, packaging and gas composition, or storage temperature will influence the outcome. Additionally, the specific microbiota of the plant may vary regionally and will engage in complex interactions with pathogens like *L. monocytogenes* (Brandl, 2006),

**Figure 3:**
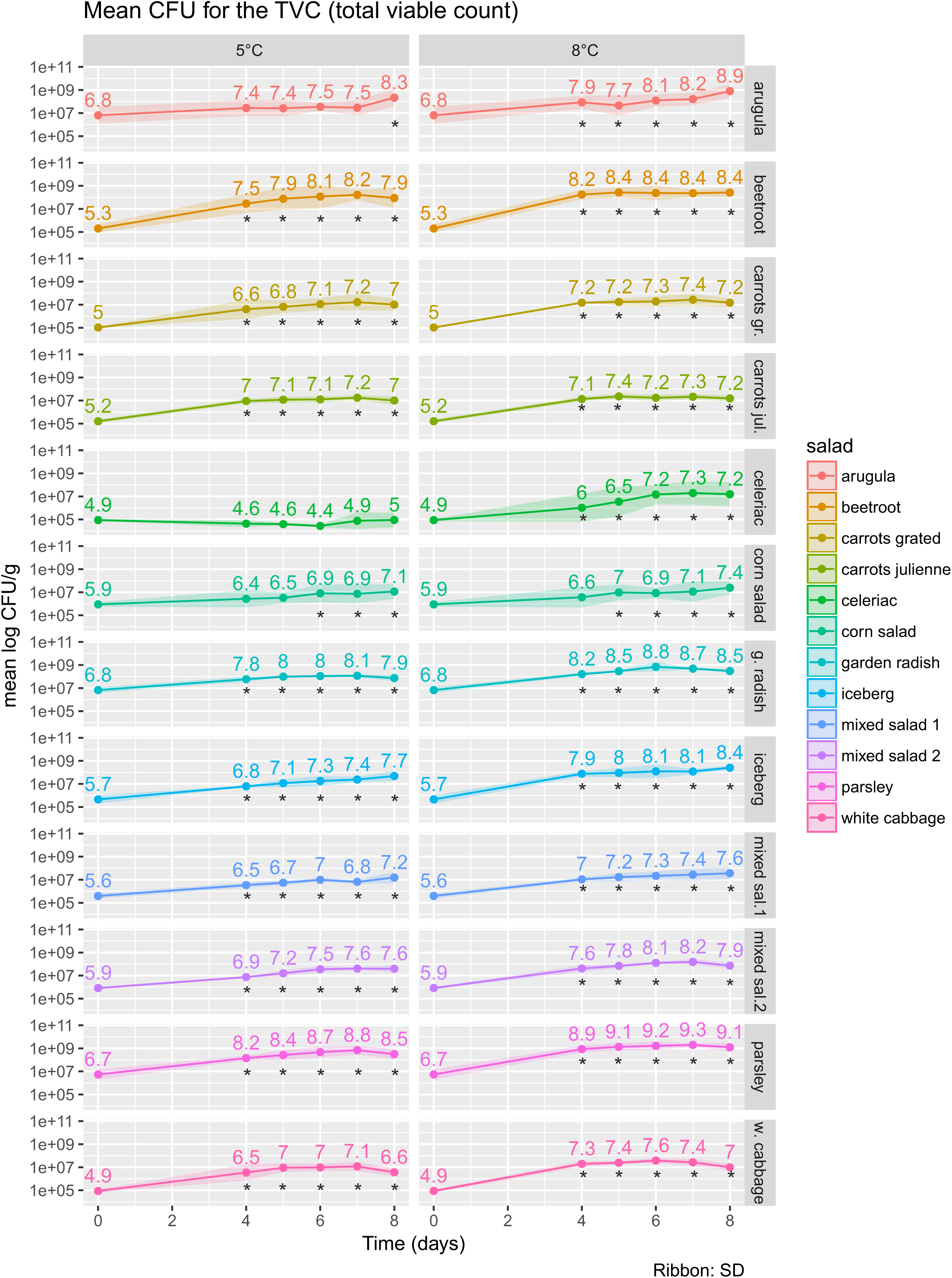
Total count of viable bacteria at 37 °C in RTE salad products. The numbers above time points reflect mean log CFU/g. Asterisks denote time points where the log CFU/g were significantly different from t=0 (p < 0.05). The ribbon around the line represents the standard deviation. gr. grated; jul. julienne; g. garden; sal. salad; w. white.

### 3.4. Total Enterobacteriaceae on RTE salad products

The initial count for Enterobacteriaceae ranged from 2.5 log CFU/g (celeriac, SD=0.61) to 5.5 log CFU/g (corn salad, SD=0.69). With the exception of corn salad, the CFU count for Enterobacteriaceae increased between t=0 and t=8 at 5 °C and at 8 °C. The sharpest increase in Enterobacteriaceae was observed in iceberg lettuce (3.5 and 3.6 log CFU/g difference between t=0 and t=8 at 5 °C and 8 °C, respectively, p < 0.001). In corn salad, a decreasing trend in Enterobacteriaceae was observed at both temperatures (−0.9 and −0.3 log CFU/g difference between t=0 and t=8 at 5 °C and 8 °C, respectively, not statistically significant) (figure 4). As most of the epiphytic microbiota of plants belongs to either *Pseudomonas* or the Enterobacteriaceae (Lund, 2008), the numbers we found are within the expected range of 4-8 log CFU/g (European Commission, Scientific Committee on Food, 2002).

**Figure 4:**
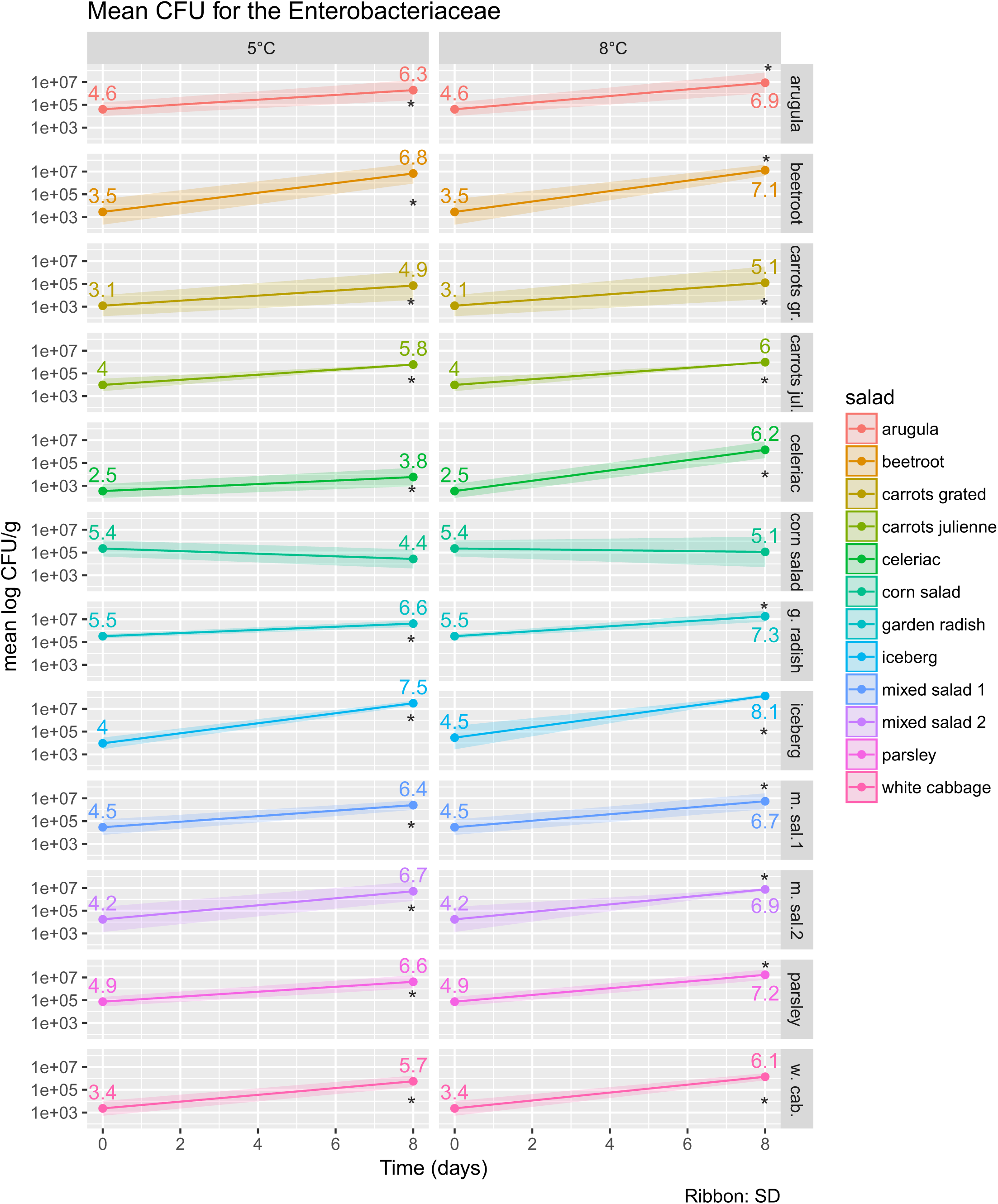
Total count of Enterobacteriaceae in RTE salad products. The numbers above time points reflect mean log CFU/g. Asterisks denote time points where the log CFU/g at t=8 were significantly different from t=0 (p < 0.05). The ribbon around the line represents the standard deviation. gr. grated; jul. julienne; g. garden; m. mixed; sal. salad; w. white; cab. cabbage.

### 3.5. Available microbial models

EC no 2073/2005 regulates the microbial criteria for foodstuffs and mandates that the food production officer shall “conduct studies (…) to investigate the compliance with the criteria during shelf-life”. Apart from challenge tests, which are labor intensive and require advanced research capacities, annex II of the EC no 2073/2005 allows the use of “predictive mathematical modelling established for the food in question” to assess the growth potential of *L. monocytogenes* in food. Existing modeling tools use physicochemical properties such as temperature, water phase salt, pH, CO2, smoke intensity, nitrite and organic acids to predict growth of microorganisms. The available models for *L. monocytogenes* are either geared towards meat / seafood (“food spoilage and safety predictor FSSP” http://fssp.food.dtu.dk, “listeria meat model” http://www.cpmf2.be/software.php, “DMRI predictive models for meat” http://dmripredict.dk) or dairy products (“dairy products safety predictor” https://aqr.maisondulait.fr). Large discrepancies were found when comparing observed growth of *L. monocytogenes* in a cheese food matrix with predictions obtained from the “Combase Modeling Toolbox” (https://www.combase.cc) in 2011, and the authors caution against applying data obtained in laboratory media to model growth in food systems (Schvartzman, Belessi, Butler, Skandamis, & Jordan, 2011). It remains a fact that the growth of pathogens on food matrix, and especially raw products, is notoriously difficult to model due to interactions with their microbiota. The linear mixed effects model calculated on the data in this study (insert link R_scripts_and_data) is perfectly valid for the dataset established in this study. However, extrapolations to other salad products would have to be done very carefully. The results are likely affected by changes in the microbiota and composition of the salad matrix due to differences in soil composition, farming practices and seasonal fluctuations in temperature and humidity. The same is true for extrapolations from public databases on microbial growth in food such as Combase (https://www.combase.cc). Therefore, even though the EU-legislation allows for the use of microbial modeling tools to assess the growth potential of *L. monocytogenes* in food, the generation of solid wet lab data in challenge tests is crucial for complex food matrices such as RTE salads composed of different raw produce.

## References

Abdul-Rauf, U. M., Beuchat, L. R., & Ammar, M. S. (1993). Survival and Growth of *Escherichia coli* 0157:H7 on Salad Vegetables. Applied and Environmental Microbiology, 59(7), 1999–2006.

Allerberger, F., & Wagner, M. (2010). Listeriosis: a resurgent foodborne infection. Clin. Microbiol Infect, 16(1), 16–23. http://doi.org/10.1111/j.1469-0691.2009.03109.x

Babic, I., Nguyen-the, C., Amiot, M. J., & Aubert, S. (2008). Antimicrobial activity of shredded carrot extracts on food-borne bacteria and yeast. Journal of Applied Bacteriology, 76(2), 135–141. http://doi.org/10.1111/j.1365-2672.1994.tb01608.x

Barton Behravesh, C., Jones, T. F., Vugia, D. J., Long, C., Marcus, R., Smith, K., et al. (2011). Deaths Associated With Bacterial Pathogens Transmitted Commonly Through Food: Foodborne Diseases Active Surveillance Network (FoodNet), 1996–2005. J. Infect. Dis, 204(2), 263–267. http://doi.org/10.1093/infdis/jir263

Beuchat, L. R., & Brackett, R. E. (1990a). Inhibitory Effects of Raw Carrots on *Listeria monocytogenes*. Applied and Environmental Microbiology, 56(6), 1734–1742.

Beuchat, L. R., & Brackett, R. E. (1990b). Survival and Growth of *Listeria monocytogenes* on Lettuce as Influenced by Shredding, Chlorine Treatment, Modified Atmosphere Packaging and Temperature. Journal of Food Science, 55(3), 755–758. http://doi.org/10.1111/j.1365-2621.1990.tb05222.x

Brandl, M. T. (2006). Fitness of human enteric pathogens on plants and implications for food safety. Annual Review of Phytopathology, 44(1), 367–392. http://doi.org/10.1146/annurev.phyto.44.070505.143359

Buchanan, R. L., Stahl, H. G., & Whiting, R. C. (1989). Effects and Interactions of Temperature, pH, Atmosphere, Sodium Chloride, and Sodium Nitrite on the Growth of *Listeria monocytogenes*. Journal of Food Protection, 52(12), 844–851. http://doi.org/10.4315/0362-028X-52.12.844

Datta, A. R., Laksanalamai, P., & Solomotis, M. (2013). Recent developments in molecular sub-typing of *Listeria monocytogenes*. Food Addit. Contam. Part a. Chem. Anal. Control Expo. Risk Assess, 30(8), 1437–1445. http://doi.org/10.1080/19440049.2012.728722

de Valk, H., Jacquet, C., Goulet, V., Vaillant, V., Perra, A., Simon, F., et al. (2005). Surveillance of listeria infections in Europe. Euro. Surveill, 10(10), 251–255.

European Commission, Scientific Committee on Food. (2002). Risk Profile on the Microbiological Contamination of Fruits and Vegetables Eaten Raw. https://ec.europa.eu/food/sites/food/files/safety/docs/sci-com_scf_out125_en.pdf

European Food Safety Authority, European Centre for Disease Prevention and Control. (2016). The European Union summary report on trends and sources of zoonoses, zoonotic agents and food-borne outbreaks in 2015. EFSA Journal, 14(12), 148. http://doi.org/10.2903/j.efsa.2016.4634

Gaul, L. K., Farag, N. H., Shim, T., Kingsley, M. A., Silk, B. J., & Hyytia-Trees, E. (2013). Hospital-acquired listeriosis outbreak caused by contaminated diced celery-Texas, 2010. Clin. Infect. Dis, 56(1), 20–26. http://doi.org/10.1093/cid/cis817

Gogo, L. A., Shitandi, A. A., Lokuruka, M. N. I., & Sang, W. (2010). Antimicrobial Effect of Juice Extract From Fermented Cabbage Against Select Food-Borne Bacterial Pathogens. Journal of Applied Sciences Research, 6, 1807–1813.

Junttila, J., Hirn, J., Hill, P., & Nurmi, E. (2016). Effect of Different Levels of Nitrite and Nitrate on the Survival of *Listeria monocytogenes* During the Manufacture of Fermented Sausage. Dx.Doi.org, 52(3), 158–161. http://doi.org/10.4315/0362-028X-52.3.158

Kaminski, C. N., Davidson, G. R., & Ryser, E. T. (2014). *Listeria monocytogenes* transfer during mechanical dicing of celery and growth during subsequent storage. Journal of Food Protection, 77(5), 765–771. http://doi.org/10.4315/0362-028X.JFP-13-382

Koseki, S., & Isobe, S. (2005). Growth of *Listeria monocytogenes* on iceberg lettuce and solid media. International Journal of Food Microbiology, 101(2), 217–225. http://doi.org/10.1016/j.ijfoodmicro.2004.11.008

Kurosaki, F., & Nishi, A. (1983). Isolation and antimicrobial activity of the phytoalexin 6- methoxymellein from cultured carrot cells. Phytochemistry, 22(3), 669–672. http://doi.org/10.1016/S0031-9422(00)86959-9

Kuznetsova, A., Bruun Brockhoff, P., & Bojesen Christensen, H. (2016). lmerTest: Tests in Linear Mixed Effects Models. Https://CRAN.R-Project.Org/Package=lmerTest.

Kühbacher, A., Cossart, P., & Pizarro-Cerdá, J. (2014). Internalization Assays for *Listeria monocytogenes*. In Cell Imaging Techniques (Vol. 1157, pp. 167–178). New York, NY: Springer New York. http://doi.org/10.1007/978-1-4939-0703-8_14

Lenth, R. V. (2016). Least-Squares Means: The R Package lsmeans. Journal of Statistical Software, 69(1), 1–33. http://doi.org/10.18637/jss.v069.i01

Lund, B. M. (2008). Ecosystems in vegetable foods. Journal of Applied Bacteriology, 73(28), 115s–126s. http://doi.org/10.1111/j.1365-2672.1992.tb03631.x

McClure, P. J., Kelly, T. M., & Roberts, T. A. (1991). The effects of temperature, pH, sodium chloride and sodium nitrite on the growth of *Listeria monocytogenes*. International Journal of Food Microbiology, 14(1), 77–91.

Popovic, I., Heron, B., & Covacin, C. (2014). Listeria: an Australian perspective (2001-2010). Foodborne Pathogens and Disease, 11(6), 425–432. http://doi.org/10.1089/fpd.2013.1697

R Core Team, R Core. (2015). R: A language and environment for statistical computing. R Foundation for Statistical Computing, Vienna.

Ragon, M., Wirth, T., Hollandt, F., Lavenir, R., Lecuit, M., Le, M. A., & Brisse, S. (2008). A new perspective on *Listeria monocytogenes* evolution. PLoS Pathogens, 4(9), e1000146. http://doi.org/10.1371/journal.ppat.1000146

R Studio Team. (2015). R Studio: Integrated Development for R. Retrieved from http://www.rstudio.com/.

Sant’Ana, A. S., Barbosa, M. S., Destro, M. T., Landgraf, M., & Franco, B. D. G. M. (2012). Growth potential of *Salmonella* spp. and *Listeria monocytogenes* in nine types of ready-to-eat vegetables stored at variable temperature conditions during shelf-life. International Journal of Food Microbiology, 157(1), 52–58. http://doi.org/10.1016/j.ijfoodmicro.2012.04.011

Schvartzman, M. S., Belessi, C., Butler, F., Skandamis, P. N., & Jordan, K. N. (2011). Effect of pH and water activity on the growth limits of *Listeria monocytogenes* in a cheese matrix at two contamination levels. Journal of Food Protection, 74(11), 1805–1813. http://doi.org/10.4315/0362-028X.JFP-11-102

Söderqvist, K. (2017). Is your lunch salad safe to eat? Occurrence of bacterial pathogens and potential for pathogen growth in pre-packed ready-to-eat mixed-ingredient salads. Infection Ecology & Epidemiology, 7(1), 1407216. http://doi.org/10.1080/20008686.2017.1407216

Stephan, R., Althaus, D., Kiefer, S., Lehner, A., Hatz, C., Schmutz, C., et al. (2015). Foodborne transmission of *Listeria monocytogenes* via ready-to-eat salad: A nationwide outbreak in Switzerland, 2013–2014. Food Control, 57, 14–17. http://doi.org/10.1016/j.foodcont.2015.03.034

Vandamm, J. P., Li, D., Harris, L. J., Schaffner, D. W., & Danyluk, M. D. (2013). Fate of *Escherichia coli* O157:H7, *Listeria monocytogenes,* and *Salmonella* on fresh-cut celery. Food Microbiology, 34(1), 151–157. http://doi.org/10.1016/j.fm.2012.11.016

Wang, J., Rahman, S. M. E., Zhao, X.-H., Forghani, F., Park, M.-S., & Oh, D.-H. (2013). Predictive Models for the Growth Kinetics of *Listeria monocytogeneson* White Cabbage. Journal of Food Safety, 33(1), 50–58. http://doi.org/10.1111/jfs.12022

Wei, H., Wolf, G., & Hammes, W. P. (2005). Combination of warm water and hydrogen peroxide to reduce the numbers of *Salmonella Typhimurium* and *Listeria innocua* on field salad (Valerianella locusta). European Food Research and Technology, 221(1-2), 180–186. http://doi.org/10.1007/s00217-005-1133-4

Werber, D., Hille, K., Frank, C., Dehnert, M., Altmann, D., Müller-Nordhorn, J., et al. (2012). Years of potential life lost for six major enteric pathogens, Germany, 2004–2008. Epidemiol. Infect, 141(05), 961–968. http://doi.org/10.1017/S0950268812001550

Wickham, H. (2009). ggplot2: elegant graphics for data analysis. In ggplot2: elegant graphics for data analysis. Springer, New York.

